# The relationship between mucosal microbiota, colitis and systemic inflammation in Chronic Granulomatous Disorder

**DOI:** 10.1101/2021.07.05.451147

**Authors:** Mehmet Davrandi, Stephanie Harris, Philip J Smith, Charles D Murray, David M Lowe

## Abstract

**Background:** Chronic granulomatous disorder (CGD) is a primary immunodeficiency which is frequently complicated by an inflammatory colitis and is associated with systemic inflammation.

**Objective:** To investigate the role of the microbiome in the pathogenesis of colitis and systemic inflammation.

**Methods:** We performed 16S rDNA sequencing on mucosal biopsy samples from each segment of 10 CGD patients’ colons, and conducted compositional and functional pathway prediction analyses.

**Results:** The microbiota in samples from colitis patients demonstrated reduced taxonomic alpha diversity compared to unaffected patients, even in apparently normal bowel segments. Functional pathway richness was similar between the colitic and non-colitic mucosa, although metabolic pathways involved in butyrate biosynthesis or utilisation were enriched in patients with colitis and correlated positively with faecal calprotectin levels. One patient with very severe colitis was dominated by *Enterococcus* spp., while among other patients *Bacteroides* spp. abundance correlated with colitis severity measured by faecal calprotectin and an endoscopic severity score. In contrast, *Blautia* abundance associated with low severity scores and mucosal health. Several taxa and functional pathways correlated with concentrations of inflammatory cytokines in blood but not with colitis severity. Notably, dividing patients into ‘High’ and ‘Low’ systemic inflammation groups demonstrated clearer separation than on the basis of colitis status in beta diversity analyses.

**Conclusion:** The microbiome is abnormal in CGD-associated colitis and altered functional characteristics probably contribute to pathogenesis. Furthermore, the relationship between the mucosal microbiome and systemic inflammation, independent of colitis status, implies that the microbiome in CGD can influence the inflammatory phenotype of the condition.

**Key Messages:** The colonic mucosal microbiome and bacterial metabolic pathways in patients with CGD colitis differ from patients without colitis, even in macroscopically normal bowel segments.

The mucosal microbiome and bacterial metabolic pathways in patients with CGD also differ according to the extent of systemic inflammation, independently from the presence of colitis, suggesting a role for the gut microbiota in the inflammatory phenotype of this condition.

**Capsule summary:** The pathogenesis of chronic granulomatous disorder (CGD)-associated colitis and other inflammatory complications is unclear. We demonstrate potentially treatable alterations in the mucosa-associated microbiome in CGD colitis and microbial differences which associate with systemic inflammation independently of colitis status.

## Introduction

Chronic granulomatous disorder (CGD) is a primary immunodeficiency characterised by failure of phagocyte oxidative burst [1]. In addition to life-threatening infection, affected patients frequently suffer inflammatory colitis [2], characterised by cryptitis, crypt abscesses and crypt architectural distortion as well as granulomas. It is known that the microbiome is altered in other inflammatory bowel diseases and that this may have a causative role in the pathogenesis [3]; this may be of especial importance in CGD where poor innate control of bacteria is a core feature. An important role for the microbiome has been suggested in a mouse model of CGD [4] and the stool microbiota has been described as abnormal in patients with CGD [5].

Microbial dysbiosis is also strongly associated with systemic inflammation of any cause, even in the absence of overt gut disease [6–8]. In a recent study, we demonstrated that CGD patients have elevated blood inflammatory markers and cytokines which do not necessarily correlate with the extent of colitis [9].

We therefore first hypothesised that the mucosa-associated microbiome and microbial metabolic pathways would differ between CGD patients with and without colitis, supporting a causative role in the development of colitis as seen in other inflammatory bowel diseases. Next, we hypothesised that the microbiome and microbial pathways would differ according to the extent of systemic inflammation regardless of the extent of colitis, implicating the microbiota in the wider inflammatory phenotype of CGD.

In our recent study, where we demonstrated that CGD-associated colitis can be monitored non-invasively [9], ten participants underwent colonoscopy with biopsies taken from each segment of the large bowel. Blood was also assayed for markers of systemic inflammation. We have here investigated the microbiota in each of these bowel segments and correlated this with the severity of the patients’ colitis and systemic inflammation.

## Methods

Patient characteristics have been described previously [9] and are provided in Table 1 and Supplementary Table E1. All patients were receiving antibiotic and antifungal prophylaxis. Colonoscopy and biopsies were performed as part of the study, although there was an urgent clinical indication in one patient with new onset (several weeks) of colitis symptoms; blood was taken contemporaneously for serum cytokine analysis.

**Table 1:** Clinical characteristics of participants in the study.

| Patient | Age (years) | Sex | CGD Type | Mutation | Total UCEIS | History of Colitis (onset) | Antibiotics | ImS |
| --- | --- | --- | --- | --- | --- | --- | --- | --- |
| **P01**<br>(nAC) | 35 | F | AR<br>(p47) | NA | 0 | Yes<br>(2014 - adulthood) | Co-trimoxazole | + |
| **P02**<br>(nAC) | 26 | M | AR<br>(p40) | NA | 0 | No | Co-trimoxazole | - |
| **P03**<br>(nAC) | 37 | M | XL | CYBB<br>c.943_945<br>delAAG,<br>p.Lys 315del | 0 | No | Co-trimoxazole | - |
| **P04**<br>(nAC) | 39 | M | XL | CYBB<br>c.252G>A,<br>splice defect | 0 | Yes<br>(unknown onset) | Co-trimoxazole | + |
| **P05**<br>(AC) | 37 | M | XL | CYBB<br>total gene<br>deletion | 7 | No* | Ciprofloxacin,<br>Doxycycline | - |
| **P06**<br>(AC) | 27 | M | XL | CYBB<br>c.665A>G,<br>p.His222Arg | 32 | Yes<br>(2006, childhood) | Co-trimoxazole | - |
| **P07**<br>(AC) | 24 | F | AR<br>(p67) | NA | 41 | Yes<br>(1997, childhood) | Co-trimoxazole | + |
| **P08**<br>(AC) | 25 | M | XL | CYBB<br>c.388C>T,<br>p.Arg130X | 9 | Yes<br>(2006, childhood) | Co-trimoxazole | - |
| **P09**<br>(AC) | 24 | M | XL | CYBB<br>deletion exon<br>6_13 | 28 | Yes<br>(2006, childhood) | Co-trimoxazole | + |
| **P10**<br>(AC) | 22 | M | XL | CYBB<br>c.676C>T,<br>p.Arg226X | 24 | Yes<br>(2015, adulthood) | Ciprofloxacin,<br>Metronidazole | + |

CGD type: XL, X-linked; AR, autosomal recessive.
ImS: Immunosuppressants.
NA: Not available.
AC, active colitis; nAC, no active colitis
\* No prior history of colitis but diagnosed with these investigations

Based on cytokine profiles, a rank scoring (1-9) was used to divide patients into two groups, (1) high level of systemic inflammation (High - below median, ranks 13-21) and (2) low level of systemic inflammation (Low - above median, ranks 29-38). Rank scores and serum IL-1β, IL-6, TNFα, IL-12 and sCD14 measurements are given in Supplementary Table E1. Serum cytokine measurements were not completed for patient P04. Colonic biopsy specimens were obtained from each bowel segment reached (rectum, sigmoid, splenic flexure, hepatic flexure, caecum, terminal ileum), and the presence of colitis in each segment was assessed by the endoscopist and scored according to the Ulcerative Colitis Endoscopic Index of Severity (UCEIS) score. Samples were stored at −80°C in RNALater (ThermoFisher).

Metagenomic DNA was extracted from approximately 1 × 1 mm biopsy sections or a blank sample (extraction control) using DNeasy PowerLyzer PowerSoil kit [11]. For 16S rDNA sequencing, V3-V4 hypervariable region of 16S rRNA gene was amplified by PCR using universal 341F and 805R primers fused with Nextera XT index and MiSeq adapter sequences. Molecular grade water was used as a negative control and Mock Community B (HM-783D, www.beiresources.org) used as a positive control, and amplicons were confirmed on a 1% agarose gel. The extraction kit control and PCR negative controls did not generate any amplicons, therefore were not included in library pooling. Subsequently, 62 samples including the mock community were pooled at equimolar concentrations and the library was sequenced using a MiSeq sequencer with 2 x 250 bp paired-end run (Illumina MiSeq, v2 kit). The resulting sequence data was processed using ‘Quantitative insights into microbial ecology 2’ (QIIME2 version 2020.2, https://qiime2.org/) [52]. The raw sequences were de-multiplexed, and de-noised using the DADA2 algorithm with default parameters to create amplicon sequence variants (ASVs). The mean sequencing depth was 63,509 (range of 9,476 – 124,026), and the resulting ASVs were assigned taxonomy using the SILVA v132 16S database. Functional metabolic predictions were calculated on ASVs using the PICRUST2 (v2.3.0-b) software with default parameters [53], and the resulting pathway functional profiles were imported into QIIME2 environment. Taxonomic profiles were generated using the 20 most abundant genera across all samples. Alpha diversity was calculated on ASVs and predicted functional pathways using the observed ASV index (number of unique features) and Shannon index. For group-wise comparisons at community level, Principal Coordinate Analysis (PCoA) was performed using the Aitchison distance [54]. The effect of active colitis, history of colitis, immunosuppression, bowel segment, age, CGD type and individuality was tested by Adonis test. The ASVs and functional pathway data sets were further standardised by analysing just two segments (sigmoid and rectum) that were available for all patients. Mann-Whitney test was used to compare the alpha diversity metrics between active colitis (AC) and no active colitis (nAC) groups as well as between high and low systemic inflammation groups. Similarly, Aitchison distances were tested for the same groupings using PERMANOVA with 999 permutations for changes in the community composition. Genus level associations for colitis status were investigated using q2-geneiss and q2-ANCOM plugins [55]. Functional pathway data were further analysed using the DEICODE plugin [56]. The resulting robust Aitchison distances were visualised using PCoA limited to 2-axes and statistical significance was tested using PERMANOVA on the basis of colitis status and systemic inflammation group (High versus Low).

Correlations between faecal calprotectin and genus level taxonomy and functional pathways were calculated using Spearman correlation on standardised data sets (i.e. two bowel segments per patient). To explore associations between gut microbiome and inflammatory markers a correlation analysis was performed between the top 20 genera and cytokine concentrations using Spearman’s rank correlation. The top 40 pathways that were associated with the separation of the high and low systemic inflammation groups on axis-1 in the PCoA were further investigated using correlation analyses with cytokine concentrations. Each of the resulting correlation matrices was clustered via hierarchical clustering using Ward’s minimum variance method (Ward.D) and Rho (r^2^) was reported. Patient P10 was excluded from some analyses, as indicated in the relevant sections, due to the extreme difference in microbial composition from other patients.

## Results

### The microbiota of CGD patients with and without colitis differs in terms of dominant taxa, alpha-diversity and beta-diversity

Consistent with existing studies, the mucosal microbiome composition showed strong inter-individuality and the differences along the bowel segments within individuals were less than the differences between individuals. Examining the dominant taxa (Figure 1A), a patient with severe acute colitis (rapid onset of symptoms over several weeks with no prior history of bowel disease) and extremely elevated faecal calprotectin (P10) had a microbiota dominated almost exclusively by *Enterococcus*. Other patients with colitis exhibited predominantly *Bacteroides* species, and in total there were nine bacterial genera which distinguished colitis patients from those without colitis (Supplementary Figure E1A). Analysis using the q2-gneiss tool revealed that increased proportions of *Bacteroides, Clostridium innocuum* group, *Escherichia – Shigella* and *Lachnoclostridium* were associated with active colitis, while greater abundance of *Blautia, Alistipes, Bifidobacterium, Dorea* and *Subdoligranulum* were associated with non-colitic gut. Notably, in colitis patients there was no clear difference between segments affected or unaffected by disease, unlike some reports in Crohn’s disease [10]. We used a secondary approach (ANCOM test) to identify differentiating genera, and the difference in abundance of *Subdoligranulum* was the only statistically significant result. The discovery of this genus - in agreement with prior studies - despite the small size of this cohort, may indicate a functional protective role against colitis development and progression [11,12].

To further investigate the drivers of the non-colitis and colitis-associated mucosal microbiome, we performed an exploratory multivariate analysis using Aitchison distances on the 16S data. The results showed significant effects of active colitis, history of colitis (HoC), age, CGD type and use of immunosuppressants (ImS), although the largest explanatory factor was patient individuality accounting for 33% of the variance (Figure 2C). The effect of active colitis was evident in 16S alpha-diversity measures in which patients with colitis (n = 5, excluding P10 with almost exclusively *Enterococcus*) demonstrated reduced taxonomic alpha-diversity compared to those without colitis (n = 4; Figure 2A). A difference in Shannon index was also seen between those with a history of colitis versus those with no prior colitis (Supplementary Figure E2) which is in agreement with trends reported previously [5]. Thus, from both alpha- and beta-diversity results, it appears that having a history of colitis is a significant underlying factor which appears to have a long-lasting effect on the composition of the gut microbiome. The use of immunosuppressants did not impact the richness and diversity of the mucosal microbiome in this cohort but did make some contribution to beta-diversity. Eight patients were receiving co-trimoxazole prophylaxis and two were on different antibiotic regimes; however, the outlier patient P10 was in the latter group and thus – although they may be an important contributing factor – analysis on the basis of antibiotics would not be informative. All patients received itraconazole as antifungal prophylaxis.

**Figure 2:**
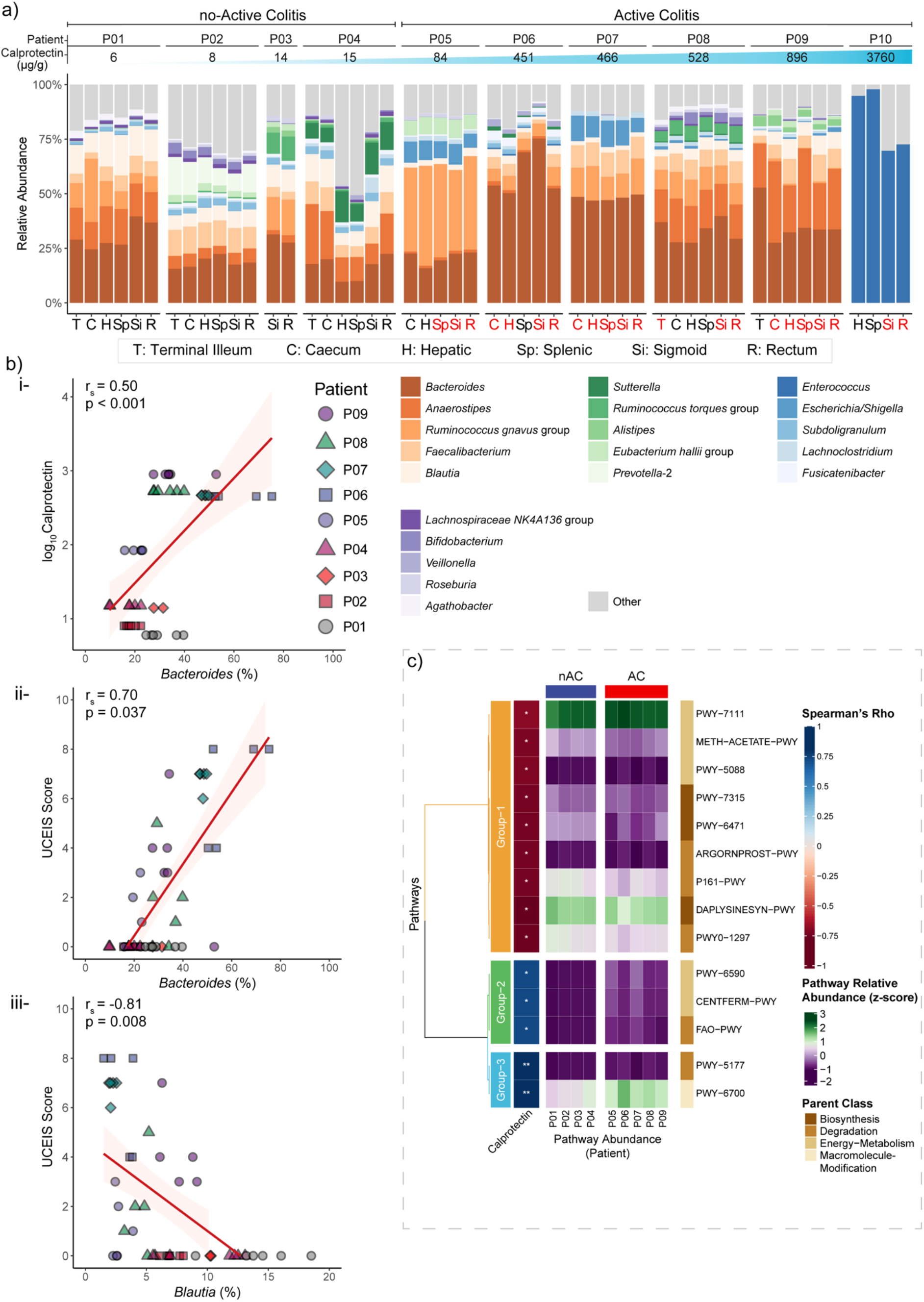
Genus level gut microbiome profiles of CGD patients with non-colitic or colitic colon. (a) Taxonomic profiles of microbiota along the gut for each patient are shown, arranged according to increasing concentrations of faecal calprotectin. In addition to overall colitis status, bowel segments with active colitis are highlighted in red. (b) Significant correlations between the faecal calprotectin level (measured per patient) and UCEIS score (measured per bowel segment) and relative abundance of *Bacteroides* and *Blautia* genus. (c) Correlation between functional pathways and faecal calprotectin level. For correlation analyses, Spearman’s correlation coefficient(r_s_) is reported. P10 was excluded from the correlation analyses because of the extreme difference in microbiome composition.

### *Bacteroides* abundance positively correlates while *Blautia* abundance negatively correlates with colitis severity

We proceeded to investigate correlations between the abundance of bacterial genera and colitis severity. Genus *Blautia* showed a strong negative correlation (r^2^ = −0.81, p = 0.008) with the endoscopic score of disease severity (UCEIS), while the genus *Bacteroides* showed a positive correlation (r^2^ = 0.70, p = 0.037) with the same measure (Figure 1B-ii,iii). Disturbances in some of these taxa have been implicated in other inflammatory bowel diseases suggesting similarities in pathogenesis, and the strong associations with these genera suggest they may be useful as indicators of colitis activity or severity in this cohort [3, 7].

In addition to the UCEIS scoring, elevated levels of faecal calprotectin are associated with intestinal inflammation which can be caused by colitis. In our patients, a level of faecal calprotectin (FCP) above 50 ug/g was indicative of active colitis, and the levels showed some correlation with the *Bacteroides* genus (r^2^ = 0.50, p < 0.001; Figure 1B-i).

### Certain microbial functional pathways correlate with colitis severity as measured by faecal calprotectin

Calprotectin is known to chelate metallic ions such as Zn, Mn, Fe and Cu, and it can therefore act as an inhibitor for metalloenzymes. This could be a potential underlying reason for the enrichment or reduction of certain metabolic capabilities which cannot be directly inferred by taxonomic assignments. To investigate the relations between the level of FCP and functional characteristics of the mucosal microbiome a correlation analysis was completed. 14 functional pathways were found to significantly correlate (p < 0.05) with the FCP level (Figure 1C). The first cluster (Group-1) contained negatively correlated pathways. Among these, DAPLYSINESYN-PWY (L-lysine biosynthesis I) pathway contains the zinc-dependent dapE metalloenzyme, which plausibly could be limited due to calprotectin mediated sequestration [13]. Moreover, this pathway has been previously identified as one of the differentially abundant pathways between healthy and ulcerative colitis patients [11].

The second cluster (Group-2) contained three functional pathways that positively correlated with the FCP levels. (1) PWY-6590, the superpathway of *Clostridium acetobutylicum* acidogenic fermentation pathway; (2) CENTFERM-PWY, the pyruvate fermentation to butanoate pathway; (3) FAO-PWY, the fatty acid β-oxidation I pathway. Notably, all three are either involved in butyrate biosynthesis or utilisation of fatty acids such as butyrate. Compared to the non-colitis patients, the butyrate producing pathways as well as butyrate utilising pathways were enriched in patients with colitis, particularly in patients P05 and P07. This suggests that reduced butyrate levels in IBD patients could be due to increased microbial utilisation of butyrate [14–16], although butyrate levels in CGD colitis have not previously been measured.

In the third cluster, which demonstrated a strong positive correlation with FCP, one of the notable pathways was the PWY-6700 (queuosine biosynthesis I) pathway. Queuosine plays an essential role in tRNA modifications, and the gut microbiome is thought to be a major provider of this micronutrient to the host [17,118]. However, it was shown to be upregulated under Zn limiting conditions, so again increased FCP may potentially be responsible for its increased abundance in active colitis patients [19,20].

### Analysis of microbial functional pathways reveals differences between CGD patients with and without colitis

Multivariate analysis using Aitchison distances on functional metabolic pathway predictions also showed significant effects of active colitis, history of colitis (HoC), age, CGD type and use of immunosuppressants (ImS). Notably, the amount of variation explained by active colitis was almost double (at 24%) that observed in the 16S sequencing data (13%). However, individuality remained the largest explanatory factor at 30% (Figure 2B-iii). In terms of alpha-diversity, only active colitis and having a history of colitis resulted in significant differences on Shannon Index (Supplementary Table E2). Although the mean difference between groups was small (~0.1) for both factors, ANCOM test revealed differentially abundant pathways. Three pathways were significantly enriched in patients with colitis. These were PWY-6590, CENTFERM-PWY and FAO-PWY, all of which also showed significant correlation with the FCP levels, as described in the next section (Supplementary Figure E1B). On the other hand, GLUCARDEG-PWY (the D-glucarate degradation I pathway) had higher abundance in patients with prior colitis.

### Systemic inflammatory markers correlate with the abundance of certain bacterial genera

We next sought to investigate the relation between blood inflammatory markers and mucosal microbiome composition. Correlation analysis showed distinct patterns clustered under four groups (Supplementary Figure E3). Group-1 mostly consisted of positively correlated genera and included the strongest significant association between *Lachnoclostridium* and IL12 (r^2^ = 0.83, p = 0.01). This genus also had increased abundance in patients with active colitis in our cohort, and has been previously shown to have increased abundance in colitis but not in Crohn’s disease [12].

Group-3 and Group-4 were almost entirely represented by negative correlations while differing in the associated inflammatory markers. The former included mucosal non-colitic *Blautia, Alistipes* and *Faecalibacterium* genera showing significant negative correlations with IL12, sCD14 and IL1β. In the latter, there were significant negative associations between the innate cytokines (IL6, IL1β and TNFα) and *Agathobacter, Bifidobacterium, Anaerostipes* and *Lachnospiraceae* NK4A136 group. The majority of these genera (*Alistipes, Faecalibacterium, Bifidobacterium* and *Lachnospiraceae* NK4A136 group) are short-chain fatty acid (SCFAs; acetate, propionate and butyrate) producing bacteria that can reduce inflammation [21]. In particular, an association between increased abundance of *Faecalibacterium prausnitzii* coupled with increased butyrate and decreased levels of sCD14 has previously been reported in HIV-infected individuals [22]. Interestingly, across the top 20 genera, we were not able to identify as many significant positive as negative correlations with the levels of inflammatory markers. In particular, despite *Bacteroides* genus showing significant correlations with the FCP levels and the endoscopic UCEIS score, it did not significantly correlate with any of the systemic inflammatory markers. This lack of positive associations also suggested that there might be additional factors other than colitis which mediate the interplay between mucosal microbiome and systemic inflammation.

### Microbial composition and functional pathway alpha-diversity metrics differ between patients with high and low levels of systemic inflammation

To explore this possibility, we introduced high (n = 4) and low (n = 4) systemic inflammation groups based on the total rank scores of the inflammatory markers (Supplementary Table E1). Notably, a patient with no colitis and one with the mildest disease were classified into the ‘High’ inflammation group while two patients with active colitis were found to have ‘Low’ ranks of inflammatory markers, implying a lack of clear correlation between colonic and systemic inflammation.

Revisiting alpha-diversity metrics, both taxonomic and functional pathway richness showed significant differences between the High and Low systemic inflammation groups (Supplementary Table E2). More importantly, inflammation was the only factor that resulted in a significant separation of the functional pathway richness. The Low systemic inflammation group had a richness mean of 282 (±3) unique functional pathways compared to 261 (±17) in the High inflammation group. Together, these results suggest either that increased systemic inflammation is reducing functional pathway richness in the gut microbiome, or conversely (and more plausibly) that reduced functional richness due to disruption of gut microbiome might be inducing inflammation.

### Multivariate analysis reveals a strong association between systemic inflammation and the gut microbiome

To further investigate the association between gut mucosal microbiome and systemic inflammation, we performed robust Aitchison PCA limited to two dimensions. The initial multivariate analyses showed active colitis as a significant factor explaining 22% and 21% of the variation in the 16S rDNA and functional pathway data, respectively (Supplementary Table E3). However, the greatest explanatory factor was systemic inflammation accounting for 37% of variation for 16S rDNA and 59% for functional pathway data, even greater than the inter-individual differences. Use of immunosuppressants, CGD type and history of colitis had more modest effects. The relationship between systemic inflammation and functional and compositional characteristics was further confirmed by PERMANOVA showing a significant and large effect size for both data types (Figure 3A,B).

**Figure 3:**
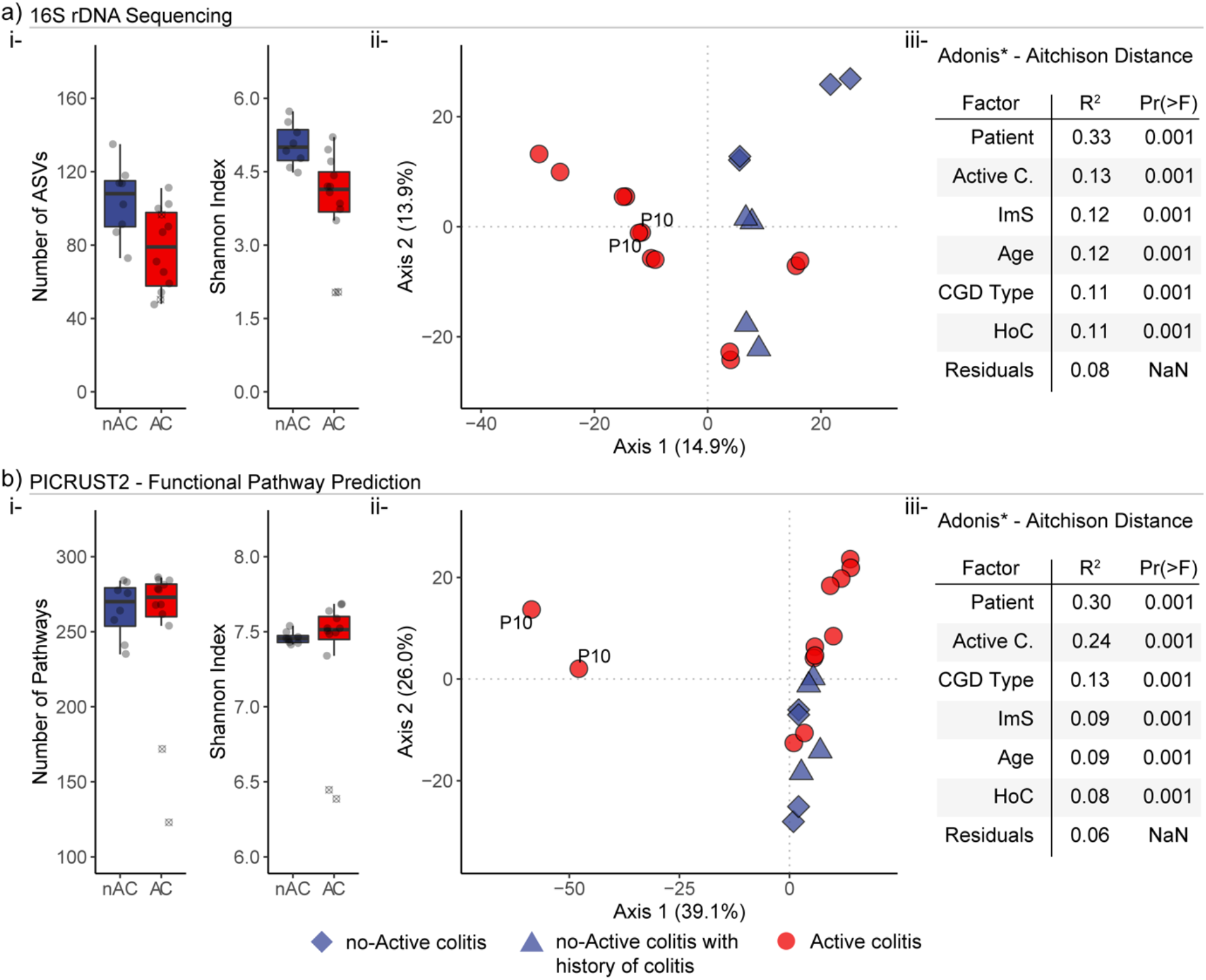
Alpha and beta diversity of the patient cohort in relation to active colitis and health. (a) based on 16S rDNA sequencing data, and (b) based on functional pathways predicted by the PICRUST2 software. (i) Richness (number of ASVs) and diversity (Shannon Index) in patients with active colitis (AC) versus those with no active colitis (nAC). Data are shown from the standardised data set with two bowel segments (rectum and sigmoid) per patient. Patient P10, who had an enterococci dominated microbiota is shown with crossed circles 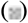. (ii) Community level clustering of the no colitis and active colitis groups; patients with a history of colitis are also indicated. (iii) Multivariate analysis by Adonis on the Aitchison distances. Adonis formula; *distance ~ Active Colitis + Immunosuppression + History of Colitis + CGD Type + Age + Gender + Patient*. Statistical significance between groups is reported in supplementary Table E2. Active colitis, n = 6; no active colitis, n = 4. Active C., active colitis; ImS, immunosuppression; HoC, history of colitis.

**Figure 3a:**
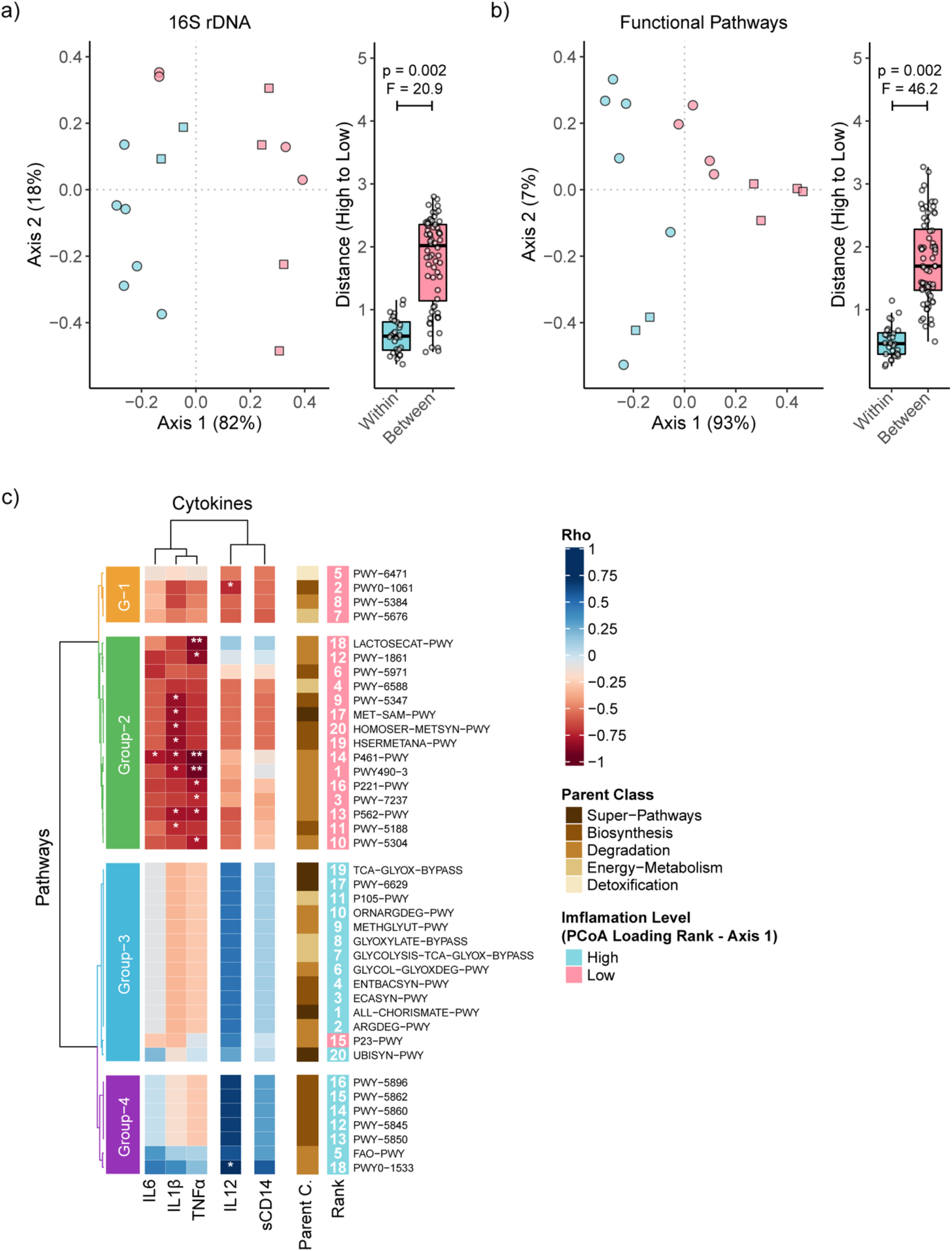
Functional pathway and taxonomic associations between mucosal microbiome and systemic inflammation. PCoA of robust Aitchison distances on the basis of systemic inflammation using (a) ASVs from 16S rDNA sequencing, and (b) functional pathway predictions. Samples with active colitis are shown as circles (O) and no active colitis as squares (□), while fill colours represent High or Low systemic inflammation. (c) Correlation analysis between the top 20 loadings (for both High and Low systemic inflammation groups) on axis 1 for functional pathways and inflammatory markers. Significance was tested by PERMANOVA as detailed in Supplementary Table E4.

Subsequently, ANCOM test revealed one genus and five pathways which were differentially abundant between the High and Low systemic inflammation groups (Supplementary Figure E4). The single genus that differed between the systemic inflammation groups was the *Lachnospiraceae* NK4A136 group, which was completely absent in the patients with High systemic inflammation. This group has been shown to be health associated in various studies [23,24], as well as considered to have anti-inflammatory properties since it is a SCFA producer [25].

We also sought to investigate whether there was any overlap between the genera associated with colitis and systemic inflammation (Figure 4). *Blautia*, *Alistipes* and *Bifidobacterium* were exclusively found to differentiate patients with both non-colitic colon and low systemic inflammation, whereas *Lachnoclostridium* was exclusively observed as a significant feature in both active colitis and high systemic inflammation. However, five further taxa differentiated colitis from non-colitis without being implicated in systemic inflammation, while six taxa were differentially abundant between the high and low inflammation groups without varying according to colitis status. Again, this implies that the relationship between gut mucosal microbiome and systemic inflammation is more complex than simply reflecting the severity of colitis.

**Figure 4:**
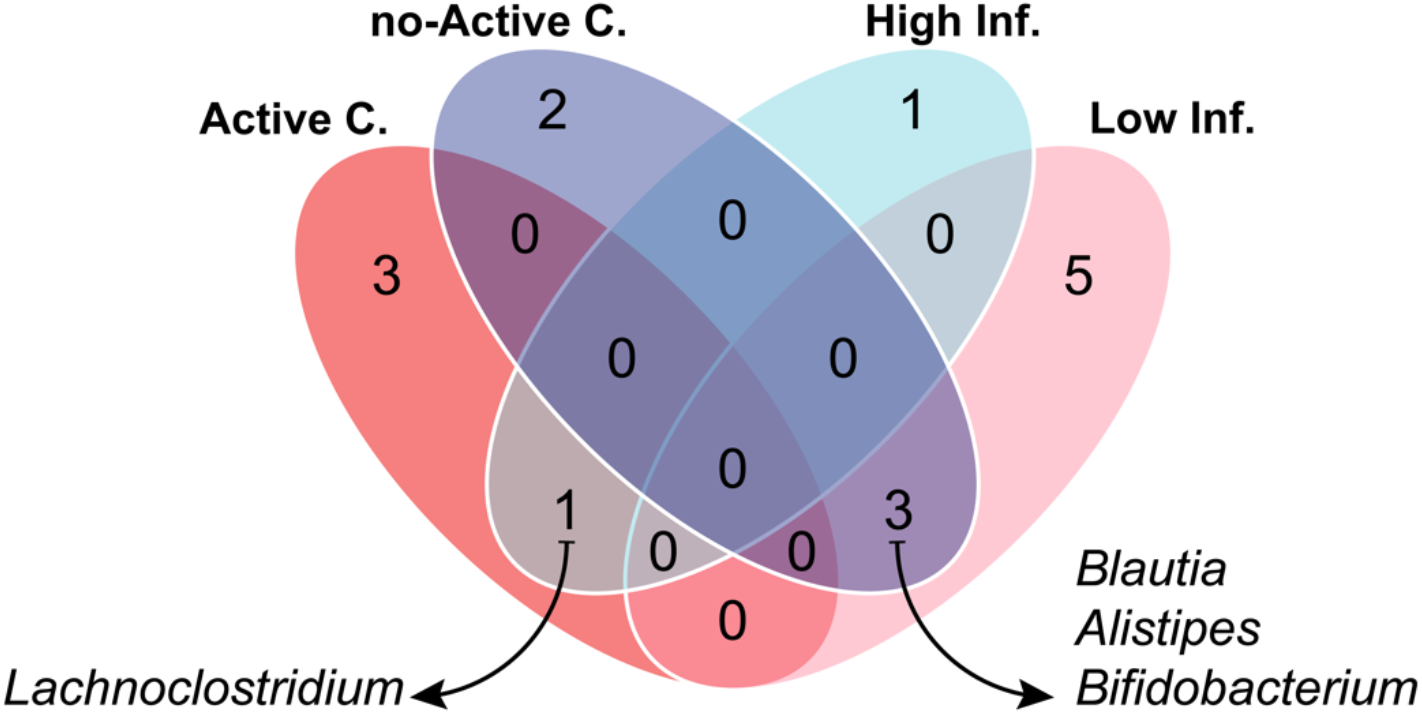
Genus level taxonomic associations shared across active colitis and systemic inflammation groupings. The Venn diagram concatenates those genera that were identified as significantly different between groups in previous analyses. *Lachnoclostridium* was shared between colitis and high levels of systemic inflammation, while *Blautia, Alistipes* and *Bifidobacterium* was associated with non-colitic colon and low level of systemic inflammation. C, colitis; Inf., inflammation.

Three of the functional pathways, P562-PWY (myo-inositol degradation I), PWY-5304 (superpathway of sulphur oxidation), P461-PWY (hexitol fermentation to lactate, formate, ethanol and acetate), had increased abundance in the Low inflammation group. Myo-inositol is abundant in the gut, and higher abundance of its degradation pathway in low systemic inflammation may reflect the presence of a non-dysbiotic mucosal microbiome. PWY-5304 is related to sulphur metabolism in which hydrogen sulphide, L-cysteine and inorganic sulphate can be produced by microorganisms such as *Desulfovibrio* species that may have immune-regulatory properties [26–28]. Lastly, P461-PWY was found to be more abundant in the Low Inflammation group. This pathway is responsible for fermentation of sugar alcohols and has been associated with *Anaerostipes hadrus* [29].

By contrast, the remaining two pathways had increased abundance in the High inflammation group, and both were involved in the biosynthesis of vitamin B6 (pyridoxal 5′-phosphate, PLP). These were PWY0-845 (superpathway of PLP biosynthesis and salvage) and PYRIDOXSYN-PWY (PLP biosynthesis I). Gut microbiota is one of the main sources of Vitamin B6, though the mechanism by which it affects host-microbiome interplay is not well established as there are conflicting reports regarding its relationship to inflammation [30–33].

### The abundance of certain microbial metabolic pathways correlates with systemic inflammatory markers

To further clarify the relation between the most implicated microbial pathways and inflammation, we performed correlation analysis between the top 20 contributors to axis-1 in the Aitchison PCA and individual cytokines (Figure 3C). Primarily, pathways that were associated with a lower level of systemic inflammation clustered in Group-1 and Group-2, and the majority of the significant interactions (14/18) occurred with IL1β and TNFα. The three pathways (PWY-562, PWY-5304, P461-PWY) that were found to have increased abundance in patients with Low systemic inflammation also showed significant negative correlations with one or more of the monocyte derived cytokines. In particular, P461-PWY negatively correlated with IL6, IL1β and TNFa, which has not been reported previously. In addition, four methionine biosynthesis pathways, PWY-5347 (superpathway of L-methionine biosynthesis), MET-SAM-PWY (superpathway of S-adenosyl-L-methionine biosynthesis), HOMOSER-METSYN (L-methionine biosynthesis I), HSERMETANA (L-methionine biosynthesis III) showed negative correlations with inflammatory markers. Similarly, a P562-PWY related pathway, PWY-7237 (myo-, chiro- and scillo-inositol degradation), showed negative correlation with TNFα.

In contrast, Group-3 and Group-4 mostly consisted of positive correlations, although there was only one significantly correlating pathway (PWY0-1533, methylphosphonate degradation I). This pathway had a strong correlation (r^2^ = 0.71) with the IL12 level, and its abundance was increased in the High inflammation group compared to the Low inflammation group. Notably, this pathway has been reported to be enriched in the dysbiotic gut microbiome of severe acute malnutrition patients with acute diarrhea [34]. It was also notable that the FAO-PWY pathway which was positively associated with increased FCP level and identified to have increased abundance in colitis, showed positive correlation (r^2^ = 0.59) with levels of IL12.

## Discussion

The current study is the first to report on the mucosal microbiome of patients with CGD in relation to colitis and systemic inflammation. Given the rarity of this condition, conducting a large scale investigation would be challenging, and to our knowledge, there is only one previous study describing the faecal microbiome of 11 patients with this immunodeficiency condition [5]. Although this previous study provided valuable first insights on the gut microbiome in CGD, other groups have demonstrated that the localised mucosal microbiota along the gut can significantly differ from that of the faecal in health as well as disease. Therefore, investigating mucosal microbiota in CGD patients may further help us in understanding the regulatory role of the gut microbiota in relation to both colitis and systemic inflammation.

As revealed by the genus level taxonomic profiles, the intra-individual differences along the gut segments were marginal in most cases. In agreement with some [35–38] but not all [10] previously reported studies, the differences between the colitic and non-colitic segments within individual patients were also minimal in terms of taxonomic composition and alpha diversity. On the other hand, the expected inter-individual difference between patients was the greatest explanatory factor in the initial beta diversity analyses independent from colitis status [39,40]. However, a number of genera were found to associate with either colitic or non-colitic gut. In particular, our results showed that elevated abundance of *Subdoligranulum* was indicative of normal gut mucosa. Although this was consistent with other studies suggesting its preventative role In IBD [11,12,41], one earlier study linked its reduced abundance in IBD patients to administration of antibiotics [42]. All patients in our study were on antibiotics, although we were unable to analyse according to the agents received as nearly all were on co-trimoxazole. The one patient with an *Enterococcus* dominated microbiome was receiving ciprofloxacin and metronidazole and we cannot exclude that these antibiotics were a contributory factor. For example, although the same pattern was not seen in the other patient on ciprofloxacin (plus doxycycline), metronidazole might theoretically have reduced *Bacteroides* abundance.

Another interesting finding was the strong negative correlation between *Blautia* and endoscopic assesment scores, in parallel with its increased abundance in non-colitis patients. This bacterium’s presumably protective role in CGD colitis contrasts with a recent study in which it was associated with the IBD-related microbial network in the gut [43]. We also observed a significant positive correlation between the *Bacteroides* genus and markers of colitis severity. Interestingly, the characteristics of the members of this genus may differ substantially, even at species level. For example, some *B. fragilis* can have anti-inflammatory and protective properties against colitis [11,44,45], while the enterotoxigenic *B. fragilis* can induce inflammation and promote IBD [46,47]. In a related study conducted by Fiedorova and colleagues [39], the authors emphasize the inconsistencies among studies in identifying certain bacteria associating with gastrointestinal disease severity or health in CVID patient cohorts. While there are several host-related (e.g. genetic makeup and CVID characteristics) and environmental factors (e.g. geographical origin, cohort size and methodology) that could impact the results of such studies, it could also imply that the taxonomic changes may not have to be consistent because the underlying driver is the total functional metabolic capability of the gut microbiota.

With this approach, i.e. analysing on the basis of microbial functional pathways, we identified an increased number of significant associations by correlation analyses and abundance testing, which improved the contribution of colitis as an explanatory factor in beta diversity analyses (albeit patient individuality still remained the main factor).

However, the lack of a clear separation between active colitis and no active colitis groups led to the introduction of systemic inflammation levels as a new variable. Surprisingly, the effect size of the existing explanatory factors (e.g. colitis status) and in particular individuality, changed drastically meaning that systemic inflammation was independent from colitis disease severity as reported previously [35]. A dysbiotic gut microbiota was shown to induce systemic inflammation in mice [48], and our findings suggest that high systemic inflammation in CGD can be associated with altered gut microbial composition and, especially, functional capability. Collectively, these results strongly implicate the colonic mucosal microbiome in the systemic inflammatory phenotype of CGD, independently of the impact of colitis.

The gut microbiome is continuously shaped by a combination of factors, and it can reveal explicit relationships in diseases such as colitis. Our findings also support the concept that there is not a single universal healthy gut microbiota composition. Host lifestyle factors, genetic background, health or specific diseases, the environment as well as aging drive microbiota composition and may result in several shifts and alterations over time [49–51]. For example, we discovered a modest but significant impact of age and genetic type of CGD (X-linked versus autosomal recessive) on beta diversity in our cohort. The latter might relate to differences in residual neutrophil function, although we have no clear evidence for this at present. Nevertheless, ultimately the microbiome should reach stable homeostasis with the host in terms of metabolic functional capability. These properties of a ‘healthy’ gut microbiome are crucial for understanding the interaction with the host as well as personalised treatment approaches, particularly in immunocompromised patients.

There are a number of limitations to our study that need to be acknowledged. Firstly, our findings are limited to the changes within a CGD patient cohort and did not include healthy individuals for comparison since recruitment of healthy patients was not possible due to sampling by colonoscopy. Secondly, the functional pathway data was generated using a computational prediction tool and may not completely reflect the true functional profiles. Lastly, non-bacterial members of the gut microbiota such as fungi may provide additional insights but were not analysed here.

## Conclusions

In this work, we have first demonstrated that patients with CGD-associated colitis exhibit reduced diversity in microbial populations at the level of the gut mucosa and identified bacterial taxa which appear to differentiate between non-colitic and colitic colon; the abundance of *Bacteroides* appears to correlate positively and *Blautia* negatively with disease activity. Very severe colitis may be associated with dominance of a single pathogenic species (e.g. *Enterococcus*). We have also demonstrated differences in microbial metabolism between patients with and without colitis and identified metabolic pathways which associate with disease severity, possibly due to an interaction with faecal calprotectin. Many of the changes in microbiota appear to persist even with mucosal healing and similar patterns are observed in both affected and unaffected segments of patients’ colons, implying a microbial ‘risk phenotype’ for the development of colitis. It will be interesting to study whether this resolves after successful haematopoietic stem cell transplant or gene therapy.

We have also demonstrated changes in microbial taxa and metabolism corresponding with systemic inflammation, which is not fully explained by the presence of colitis. Indeed, inflammation appears to show clearer and more significant associations with the gut microbiota than colitis itself. Our data therefore imply that CGD patients’ microbiome may influence the inflammatory phenotype of this disease and this demands further investigation. If confirmed in other cohorts, strategies to modify the gut microbiome (including faecal transplant) should be explored as therapies in CGD.

## Abbreviations

AC: Active colitis
ANCOM: Analysis of composition of microbes
ASVs: Amplicon sequence variants
CGD: Chronic granulomatous disorder
CVID: Common variable immune deficiency
FCP: Faecal calprotectin
HoC: History of colitis
IBD: Inflammatory bowel disease
ImS: Immunosuppressants
nAC: no Active colitis
PCA: Principal component analysis
PCoA: Principal coordinate analysis
PERMANOVA: Permutational multivariate analysis of variance
PLP: Pyridoxal 5’-phosphate
SCFAs: Short chain fatty acids
UCEIS: Ulcerative colitis endoscopic index of severity

## Acknowledgements

Not applicable.

## Declarations

### Availability of data and material

The datasets generated and analysed during the current study are available in the NCBI Sequence Read Archive (http://www.ncbi.nlm.nih.gov/sra) at NCBI BioProject ID: PRJNA613382

### Code availability

Not applicable

### Ethics approval and consent to participate

All patients provided written informed consent (NHS Research Ethics Committee (REC) 15/LO/1334).

### Consent for publication

All participants consented for the results of the study to be published. No individual details, images or videos are included in this manuscript.

## Authors’ contributions

MD performed laboratory and bioinformatic analyses. SH collected clinical data. PJS and CDM helped to conceive the study, performed endoscopy with assessment of colitis activity and obtained biopsies. DML conceived the study, recruited patients and supervised the analysis. All authors read and approved the final manuscript.

