## Supplemental Data for "The relationship between mucosal microbiota, colitis and systemic inflammation in Chronic Granulomatous Disorder"

### Supplementary Tables

**Supplementary Table E1: Summary of serum cytokine profiles and patient characteristics.**

| Patient | IL1β pg/mL | IL6 pg/mL | IL12 pg/mL | TNFα pg/mL | sCD14 mcg/mL | Total Rank Score (Group) | Acne / Folliculitis | Other Dermatological Diagnoses | Brch. | Inflamm. Lung Changes[1] | Emph. |
|---|---|---|---|---|---|---|---|---|---|---|---|
| **P01** | 8.3 (6) | 61 (6) | 25 (5) | 0 (9) | 6.46 (3) | 29 (Low) | Yes | Orofacial granulomatosis, eczema | No | Yes | No |
| **P02** | 4.8 (8) | 22 (9) | 3.5 (8) | 2.2 (5) | 4.22 (8) | 38 (Low) | Yes | Seborrheic dermatitis, hyperplastic granulomatous inflammation | No | No | No |
| **P03** | 40 (2) | 767 (1) | 0 (9) | 13 (1) | 5.42 (6) | 19 (High) | Yes | Hidradenitis | No | No | Yes |
| **P04** | NA | NA | NA | NA | NA | - | Yes | Nil | No | Yes | Yes |
| **P05** | 13 (5) | 74 (5) | 45 (4) | 2.0 (6) | 6.69 (1) | 21 (High) | No | Nil | Yes | Yes | No |
| **P06** | 60 (1) | 323 (4) | 132 (2) | 3.0 (4) | 6.51 (2) | 13 (High) | Yes | Nil | No | Yes | No |
| **P07** | 40 (3) | 528 (2) | 185 (1) | 6.2 (3) | 5.57 (5) | 14 (High) | No | Nil | No | Yes | Yes |
| **P08** | 5.3 (7) | 50 (8) | 24 (6) | 1.3 (7) | 6.17 (4) | 32 (Low) | No | Eczema | Yes | Yes | No |
| **P09** | 3.8 (9) | 55 (7) | 14 (7) | 0.4 (8) | 5.38 (7) | 38 (Low) | Yes | Nil | Yes | Yes | No |
| **P10** | 25 (4) | 350 (3) | 76 (3) | 6.2 (2) | 2.94 (9) | 21 (High) | Yes | Nil | No | Yes | Yes |

Brch., Bronchiectasis; Emph., Emphysema. [1] Ground glass, consolidation or scarring.

**Supplementary Table E2: Statistical significance of alpha- and beta-diversity metrics in 16S rDNA sequencing and functional pathway prediction data per grouping.**

| | Factor | Groups | N* | Richness p-value[a] | Diversity p-value[a] | Distances p-value[b] | Pseudo-F |
|---|---|---|---|---|---|---|---|
| **16S rDNA Sequencing** | Active Colitis | Yes | 5 | 0.021 | 0.001 | 0.002 | 2.30 |
| | | No | 4 | | | | |
| | History of Colitis | Yes | 7 | 0.055 | 0.015 | 0.001 | 2.47 |
| | | No | 2 | | | | |
| | Immunosuppressants | Yes | 4 | 0.625 | 0.930 | 0.002 | 2.20 |
| | | No | 5 | | | | |
| | Systemic Inflammation | High | 4 | 0.028 | 0.141 | NA | NA |
| | | Low | 4 | | | | |
| | CGD Type | XL | 6 | 0.925 | 0.639 | 0.007 | 1.88 |
| | | AR | 3 | | | | |
| | **Factor** | **Groups** | **N*** | **p-value[a]** | **p-value[a]** | **p-value[b]** | **Pseudo-F** |
| **Functional Pathways** | Active Colitis | Yes | 5 | 0.265 | 0.026 | 0.003 | 5.14 |
| | | No | 4 | | | | |
| | History of Colitis | Yes | 7 | 0.122 | 0.044 | 0.009 | 3.7 |
| | | No | 2 | | | | |
| | Immunosuppressants | Yes | 4 | 0.721 | 0.213 | 0.213 | 1.31 |
| | | No | 5 | | | | |
| | Systemic Inflammation | High | 4 | 0.005 | 0.462 | NA | NA |
| | | Low | 4 | | | | |
| | CGD Type | XL | 6 | 0.425 | 0.779 | 0.196 | 1.45 |
| | | AR | 3 | | | | |

Alpha diversity was tested on richness (number of ASVs/Pathways) and diversity (Shannon Index).
Beta diversity was tested on Aitchison distances. Inflammatory marker data was not available for P04, therefore they were excluded from the systemic inflammation grouping.
*, excluding P10; [a] statistical significance was tested by Kruskal-Wallis test; [b] statistical significance was determined by PERMANOVA with 999 permutations.

**Supplementary Table E3: Multivariate analysis (Adonis test) on robust Aitchison distances.**

Excluding P04 and P10. Adonis formula; *distance ~ Active Colitis + Immunosuppression + History of Colitis + Systemic Inflammation + CGD Type + Patient.*

| | **16S rRNA Sequencing** | | **Functional Pathway Predictions** | |
|---|---|---|---|---|
| **Factor** | $R^2$ | **Pr(>F)** | $R^2$ | **Pr(>F)** |
| Active Colitis | 0.22 | 0.001 | 0.21 | 0.002 |
| Immunosuppressants | 0.16 | 0.001 | 0.02 | 0.090 |
| CGD Type | 0.05 | 0.001 | 0.08 | 0.007 |
| History of Colitis | 0.02 | 0.039 | 0.02 | 0.080 |
| Systemic Inflammation | 0.37 | 0.001 | 0.59 | 0.001 |
| Patient | 0.15 | 0.002 | 0.03 | 0.109 |
| Residuals | 0.03 | NaN | 0.05 | NaN |

**Supplementary Table E4: Group-wise comparisons (PERMANOVA) on robust Aitchison distances.**

| | | | **16S rRNA Sequencing** | | **Functional Pathway Predictions** | |
|---|---|---|---|---|---|---|
| **Factor** | **Groups** | **N** | **p-value** | **Pseudo-F** | **p-value** | **Pseudo-F** |
| Active Colitis | Yes | 5 | 0.050 | 4.1 | 0.071 | 3.6 |
| | No | 3 | | | | |
| History of Colitis | Yes | 6 | 0.314 | 0.9 | 0.464 | 0.6 |
| | No | 2 | | | | |
| Immunosuppressants | Yes | 3 | 0.147 | 2.2 | 0.684 | 0.2 |
| | No | 5 | | | | |
| Systemic Inflammation | High | 4 | 0.002 | 20.9 | 0.002 | 46.2 |
| | Low | 4 | | | | |
| CGD Type | XL | 6 | 0.023 | 1.51 | 0.061 | 4.32 |
| | AR | 3 | | | | |

### Supplementary Figures

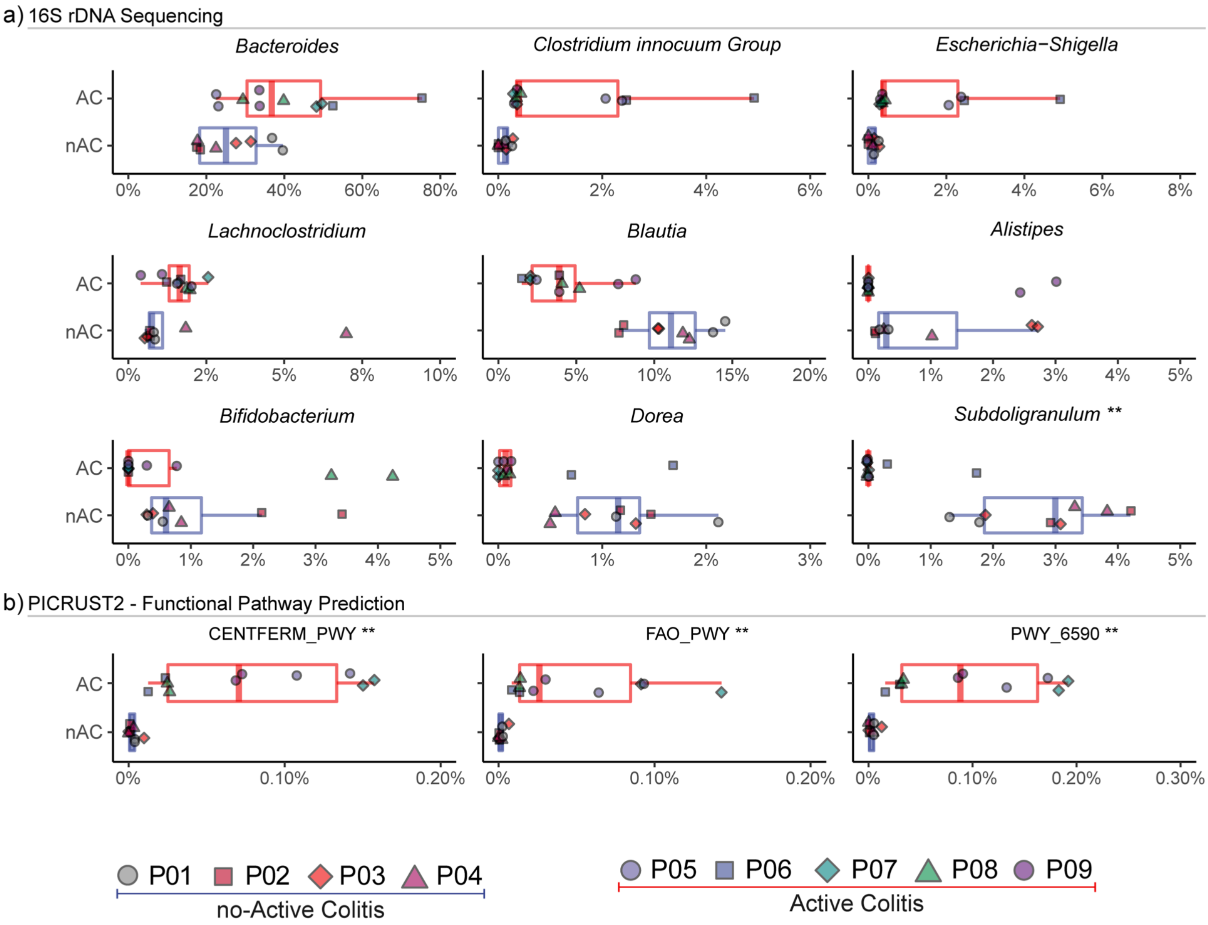


**Supplementary Figure E1: Differentially abundant genera and functional pathways among patients with active colitis.** Boxplots showing the relative abundance of genera (a) and functional pathways (b) which differ between patients without colitis (no active colitis, nAC) and those with active colitis (AC) based on the standardised data set (two segments per patient); Patient P10 was excluded from this analysis. Genus level associations were determined by the q2-gneiss tool, and further statistical testing on genus and functional pathway data was done by the ANCOM test (** denotes significance at $p \leq 0.05$). CENTFERM-PWY, pyruvate fermentation to butanoate; FAO-PWY, fatty acid and beta-oxidation I; PWY-6590, superpathway of Clostridium acetobutylicum acidogenic fermentation.

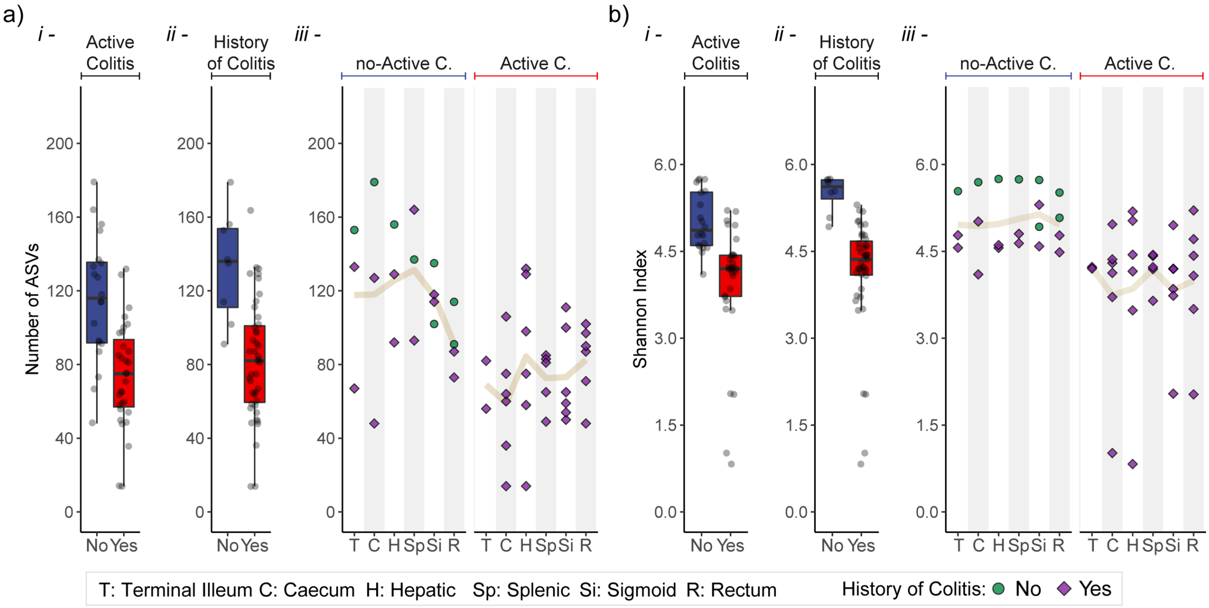


**Supplementary Figure E2: Effect of history of colitis on richness and diversity along the bowel.** Although currently having normal bowel mucosa, patients who had a history of colitis appeared to have reduced richness measured by number of ASVs (a) and diversity measured by Shannon index (b) in comparison to patients who have never experienced colitis. (i) shows the overall difference between the active colitis and no active colitis groups and (ii) shows the effect of having a history of colitis, while (iii) shows the alpha diversity landscape along the bowel. The data shown here are from all available bowel segments for each of the patients, including patient P10.

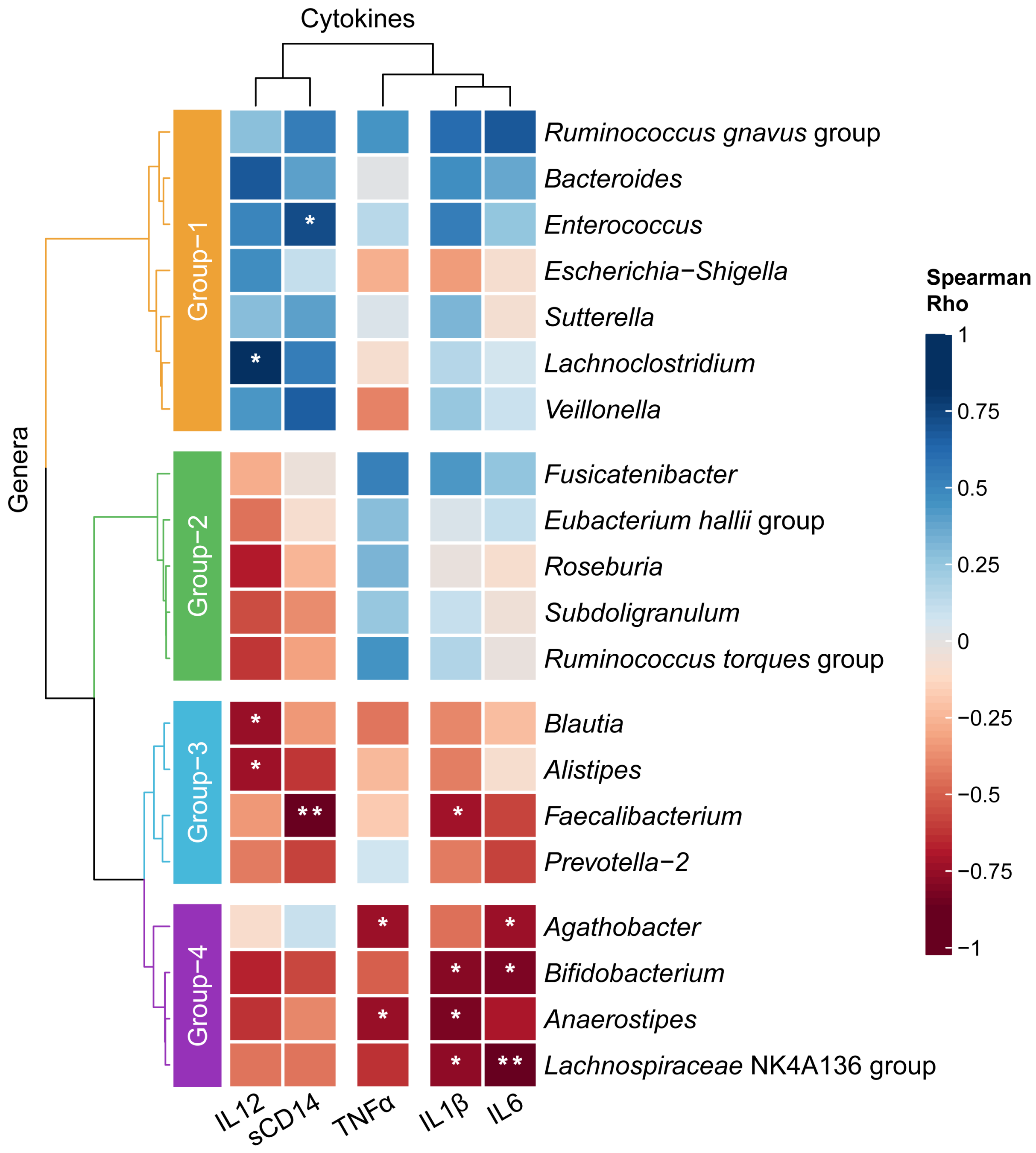


**Supplementary Figure E3: Correlation analysis between the top 20 genera and inflammatory markers.** The average of two bowel segments (rectum and sigmoid) per patient was used to calculate correlations. P10 was excluded.

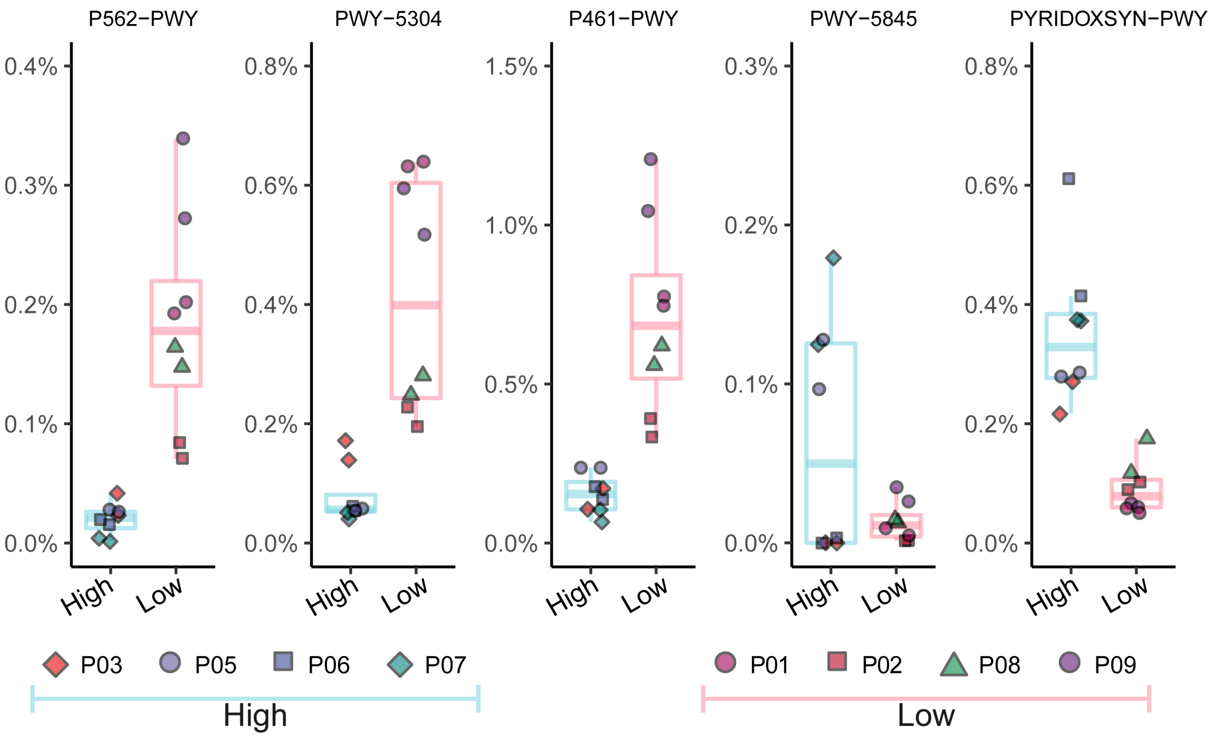


**Supplementary Figure E4: Differentially abundant functional pathways between patients with High and Low systemic inflammation.** Boxplots displaying the relative abundance of functional pathways that showed significant differences between the high and low systemic inflammation groups. ANCOM test was used to determine differentially abundant functional pathways, and a total of five pathways demonstrated a statistical significance at $p \leq 0.05$ level. P562-PWY, myo-inositol degradation I; PWY-5304, superpathway of sulfur oxidation; P461-PWY, hexitol fermentation to lactate, formate, ethanol and acetate; PWY-5845, superpathway of menaquinol-9 biosynthesis; PYRIDOXSYN-PWY, pyridoxal 5'-phosphate biosynthesis I.
